# HIV-1 Infection Regulates Gene Expression by Altering Alternative Polyadenylation Through CPSF6 and CPSF5 Delocalization

**DOI:** 10.1101/2025.08.07.669137

**Authors:** Charlotte Luchsinger, Annie Zhi Dai, Hari Yalamanchili, Aiswarya Balakrishnan, Kai-Lieh Huang, Cinzia Bertelli, Bin Cui, Ramon Lorenzo-Redondo, Eric J. Wagner, Felipe Diaz-Griffero

## Abstract

HIV-1 viral core transport to the nucleus, an early infection event, triggers cleavage and polyadenylation specificity factor (CPSF)5 and CPSF6 to translocate from paraspeckles to nuclear speckles, forming puncta-like structures. CPSF5 and CPSF6 regulate alternative polyadenylation (APA), which governs approximately 70% of gene expression. APA alters the lengths of mRNA 3’-untranslated regions (3’-UTRs), which contain regulatory signals influencing RNA stability, localization, and function. We investigated whether HIV-1 infection–induced changes in CPSF5 and CPSF6 subcellular localization are accompanied by changes in cellular function. Using two independent methodologies to assess APA in human primary CD4+ T cells and cell lines, we found that HIV-1 infection regulates APA, dependent on the interaction of CPSF6 with the viral capsid, recapitulating the APA phenotype observed in CPSF6 knockout cells. Our study demonstrates that HIV-1 infection leverages the interaction between the viral capsid and CPSF6 to co-opt cellular processes, alter gene expression, and drive pathogenesis.

## INTRODUCTION

The delivery of the HIV-1 viral core into the cytoplasm of human cells during infection requires the fusion of the viral membrane with the cellular membrane. The viral core comprises 1500 monomers of the p24 capsid protein and houses the viral RNA genome. Upon delivery to the cytoplasm, the viral core travels to the nucleus, where viral RNA is converted into viral DNA by reverse transcription ^1–3^ and the viral core undergoes uncoating, during which the monomeric capsid proteins are dissociated from the core, opening the structure housing the viral genome. Upon HIV-1 infection, and simultaneous with these early replication events, the cleavage and polyadenylation specificity factor (CPSF)5 and CPSF6 are recruited to nuclear speckles from paraspeckles ^2,4–7^. Nuclear speckles are distinct, dynamic, membrane-less structures found in the nucleus ^8,9^.

The role of human CPSF6 in HIV-1 replication has been extensively studied in both cell lines and human primary cells, revealing that HIV-1 infectivity is mildly affected by either the partial or complete depletion of CPSF6 expression ^10–12^. However, several alternative roles for CPSF6 during HIV-1 infection have been proposed, including (1) facilitating HIV-1 core entry into the nuclear compartment ^13–17^, (2) influencing HIV-1 integration and integration site selection ^11,16,18–20^, and (3) assisting with HIV-1 disassembly ^21^.

HIV-1 infection alters the immunofluorescence microscopy patterns of CPSF5 and CPSF6 from diffuse nuclear staining to easily recognizable large condensates or puncta-like structures ^2,4,5,14^ that colocalize with the nuclear speckle marker SC35, implying that HIV-1 infection triggers the translocation of CPSF5 and CPSF6 to bona fide nuclear speckles ^2,4,5,22^. However, viruses with N74D, A77V, or N57S mutations in the capsid protein fail to induce this translocation, indicating that this event is dependent on the capsid protein ^2,4,5^. Although the translocation of CPSF5 and CPSF6 to nuclear speckles is widely accepted, the effects of this translocation on the cellular functions of CPSF5 and CPSF6 remain unknown.

CPSF5 and CPSF6 are essential components of the cleavage factor I mammalian (CFIm) complex, which plays a critical role in selecting the appropriate polyadenylation signal during mRNA processing ^23,24^. Most genes contain more than one polyadenylation signal, suggesting that mRNAs may be regulated by alternative polyadenylation (APA), an essential mechanism that controls approximately 70% of human gene expression ^25^. The CFIm complex promotes the selection of distal rather than proximal polyadenylation signals, resulting in longer 3’ untranslated regions (3’UTRs) ^24^. The 3’UTR contains regulatory elements that influence key mRNA metabolic properties, including mRNA stability, translation efficiency, nuclear export, and cellular localization ^25–27^. When CFIm components are depleted, or CFIm activity is disrupted, proximal polyadenylation signals become more commonly selected, leading to shorter 3’UTRs ^28,29^ ^30^. For example, the specific depletion of CPSF5 or CPSF6 in human cells leads to the shortening of mRNA 3’UTRs ^31–33^.

The functional consequences of HIV-1 infection–induced CPSF5 and CPSF6 translocation to nuclear speckles are not understood. Because CPSF5 and CPSF6 control APA, we tested the hypothesis that HIV-1 infection modulates the APA functions of CPSF5 and CPSF6. We infected human primary CD4^+^ T cells and cell lines with HIV-1 and used two independent methodologies to measure changes in APA. We found that the infection of human primary T cells and cell lines with wild-type HIV-1 triggered APA changes, whereas APA changes were not induced by infection with HIV-1 bearing N74D or A77V mutation in the capsid protein, which prevents the capsid protein from interacting with CPSF6. Remarkably, human primary T cells infected with wild-type HIV-1 showed an increase in transcripts with shorter 3’UTRs, which ultimately altered global cellular gene expression. To model these results, we utilized the human cell line A549, a cellular model in which HIV-1 can trigger CPSF5 and CPSF6 translocation to nuclear speckles in 80-90% of infected cells at a multiplicity of infection (MOI) of 2. APA analysis revealed changes in the polyadenylation signal used by a number of genes, resulting in an increase in the number of transcripts with shorter 3’UTRs. However, infection with HIV-1 bearing the capsid mutants N74D and A77V did not induce APA changes, suggesting that HIV-1 infection–induced APA changes require the direct interaction of the capsid protein with CPSF6. Unlike HIV-1 bearing the capsid mutations N74D or A77V, wild-type HIV-1 induced the translocation of CPSF5 and CPSF6 to nuclear speckles, suggesting that the ability of HIV-1 to modulate cellular APA depends on the interaction of capsid with CPSF6. Overall, our study demonstrates that HIV-1 infection exerts control over cellular gene expression by leveraging the interaction between the viral capsid and CPSF6 to alter the subcellular localization of CPSF5 and CPSF6, impacting their functions.

## RESULTS

### HIV-1 infection induces significant changes in alternative polyadenylation

In human cells, HIV-1 infection triggers the translocation of CPSF5 and CPSF6 to nuclear speckles; however, the efficiency of this process varies across cell lines ^34^. In the human A549 cell line, HIV-1 infection at an MOI of 2 is sufficient to induce CPSF5 and CPSF6 translocation in 80% of cells ^34^, whereas translocation is only observed in 20% of HeLa cells and 1% of THP-1 cells at an MOI of 2 ^34^. Due to the high efficiency of HIV-1 infection–induced CPSF5 and CPSF6 translocation to nuclear speckles in human A549 cells, we initially studied HIV-1-infection–induced changes in cellular expression in this cell line.

We infected A549 cells with an HIV-1 construct modified to express green fluorescent protein (HIV-1-GFP) as a reporter of infection at an MOI of 2. After 24 h, we assessed infection success as the percentage of GFP-positive cells (Figure 1A) and evaluated CPSF6 translocation by quantifying the percentage of infected cells containing CPSF6 colocalized with the known nuclear speckle marker SC35 (Figure 1B), finding that 80%–90% of infected cells contained CPSF6 in nuclear speckles (Figure 1C), whereas mock-infected cells did not display CPSF6 in nuclear speckles.

**Figure 1.**
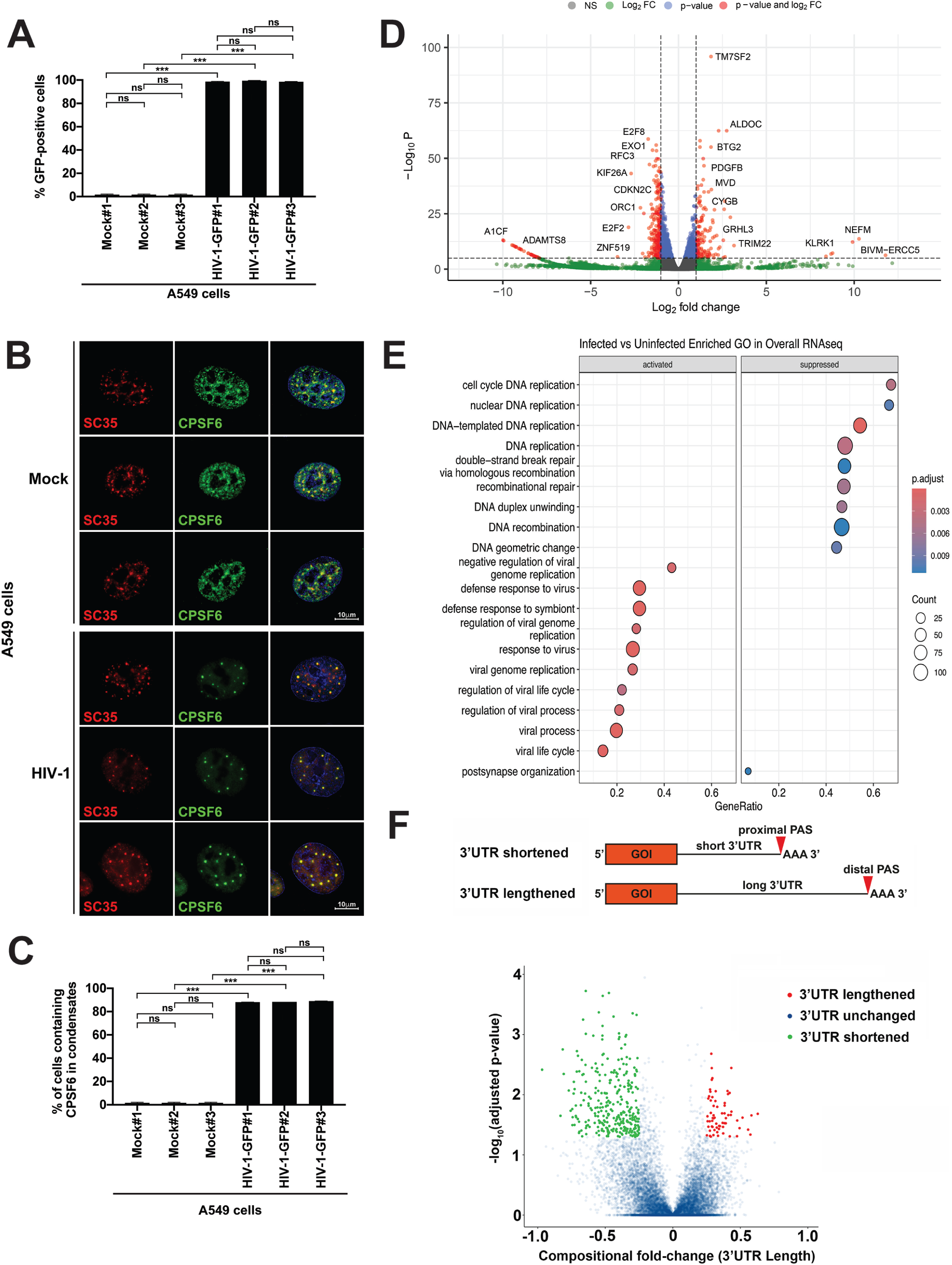
HIV-1 infection induces changes in APA. **(A)** To induce the translocation of CPSF6 to nuclear speckles, human A549 cells were challenged with HIV-1-GFP at an MOI of 2 for 24 h. Infection was assessed as the percentage of GFP-positive cells by flow cytometry. The results of three independent experiments with standard deviation are shown. **(B)** Cells were subsequently fixed, permeabilized, and immunolabelled using specific antibodies against SC35 (red, marker for nuclear speckles) and CPSF6 (green). Nuclei were stained with DAPI (blue). **(C)** The average (and standard deviation) percentage of cells containing CPSF6 in nuclear speckles (SC-35 positive compartments) from three independent experiments, as determined by visual examination of 200 cells. **(D)** Total RNA from three infected and three mock-infected samples was sequenced, and gene expression is displayed in a Volcano plot. The x-axis shows Log_2_ fold changes in gene expression, with positive indicating upregulation and negative indicating downregulation. The y-axis shows the statistical significance, expressed as −Log_10_P. **(E)** Gene ontology analysis of all genes upregulated and downregulated by HIV-1 infection. The number of genes in each pathway is expressed as the size of the sphere, and the color represents the P-adjusted value. **(F)** Volcano plot showing PAS changes. The x-axis shows compositional fold-change, which represents the 3’UTR length for each transcript. A positive compositional fold-change indicates the presence of a longer 3’UTR (red), and a negative compositional fold-change indicates the presence of a shorter 3’UTR (green). The y-axis shows the statistical significance of the result conveyed as the −Log_10_P. APA, alternative polyadenylation; CPSF6, cleavage and polyadenylation specificity factor subset 6; DAPI, 4’6-diamino-phenylindole; GFP, green fluorescent protein; MOI, multiplicity of infection; PAS, polyadenylation signal; UTR, untranslated region.

Given extensive prior reports indicating roles for CPSF5 and CPSF6 in pre-mRNA cleavage and APA, the dramatic change in subcellular localization of these factors in response to HIV-1 infection suggests that viral infection could impact polyadenylation site selection, ultimately leading to altered gene expression ^35^.

To evaluate whether HIV-1 infection changes gene expression, we obtained total RNA from three HIV-1–infected and three mock-infected A549 samples and performed RNA-sequencing to determine the presence and quantity of RNA molecules (Figure 1D–F). Changes in gene expression triggered by HIV-1 infection were plotted on a volcano plot, with the magnitude (as Log_2_ fold-change) of change plotted on the x-axis (positive values, increased expression; negative values, decreased expression), and the p-value (as −Log_10_P, with higher values indicating more significant changes) plotted on the y-axis (Figure 1D). HIV-1 infection of A549 cells resulted in the significant upregulation of 334 genes and the significant downregulation of 4589 genes, indicating that HIV-1 infection dramatically changes the cellular gene expression profile.

Our studies showed that HIV-1 infection dramatically altered cellular gene expression, indicating that HIV-1 may utilize specific mechanisms. Gene ontology analysis of activated and suppressed pathways (Figure 1E) showed that HIV-1 infection activated genes involved in countering viral infection (response to virus, viral processes, and others) and suppressed genes involved in DNA metabolism (DNA replication, recombination, and repair). Gene expression analysis using the Reactome database revealed that HIV-1 infection activated genes involved in countering viral infection (Interferon [IFN]α/B. IFNγ, IFN signaling, and others) and suppressed genes involved in cell division (M phase, mitotic anaphase, and cell cycle checkpoints) (Figure S1A).

We and others have described the translocation of CPSF6 to nuclear speckles^2,4,5^; however, the effect this translocation on CPSF6 function is not understood. This translocation to nuclear speckles may inhibit the cellular functions of CPSF6, including APA regulation. APA is a major gene expression regulatory mechanism, with 70% of human genes containing proximal and distal APA signs in the 3’UTR ^25^. Because CPSF5 or CPSF6 depletion is known to induce the use of proximal polyadenylation signals ^11,14,31,32,36–38^, we tested the hypothesis that the translocation of CPSF5 and CPSF6 to nuclear speckles during HIV-1 infection affects their functions and leads to APA dysregulation ^35^. We used regression of polyadenylation compositions (REPAC) to analyze APA in total RNA-sequencing data from HIV-1–infected and mock-infected A549 cells ^39^. We identified 246 genes associated with shorter 3’UTRs and 65 genes associated with longer 3’UTRs in HIV-1–infected cells than in mock-infected cells (Figure 1F). These results showed significant 3’UTR shortening in HIV-1–infected cells compared with mock-infected cells, suggesting that HIV-1 infection induces 3’UTR shortening and the use of proximal polyadenylation sites, phenotypically resembling the APA patterns observed with the knockout (KO) of CFIm complex components, such as CPSF5 and CPSF6. These findings suggest that HIV-1 infection interferes with the functions of CPSF5 and CPSF6.

To determine whether changes in the 3’UTR length affect gene expression, we analyzed changes in the expression of genes with HIV-1 infection–induced changes in 3’UTR length (Figure 1F). Of the 246 genes with shortened 3’UTRs following HIV-1 infection, 87 demonstrated altered transcript levels, including 82 with reduced transcript levels; however, APA may alter protein expression without altering transcript levels, as the 3’UTR can impact translation efficiency and transcript localization. By contrast, 47 of the 65 genes with longer 3’UTRs following HIV-1 infection showed altered transcript levels, including 45 with reduced transcript levels. These findings indicate that HIV-1 infection–induced changes in APA alter cellular gene expression.

We used Reactome to analyze the cellular pathways associated with genes modulated by HIV-1 infection and found that most of the genes downregulated by HIV-1 infection–induced changes in APA control cellular process, such as SUMOylation, organelle biogenesis, transport, and general transcription (Figure S1B).

### HIV-1 infection–induced changes in alternative polyadenylation are dependent on the translocation of CPSF5 and CPSF6 to nuclear speckles

Capsid proteins bearing the N74D and A77V mutations bind CPSF6 at significantly lower levels than wild-type capsid proteins ^40^. To investigate the role played by HIV-1 infection–induced CPSF5 and CPSF6 translocation in infection-induced changes in APA, we infected A549 cells with HIV-1 bearing the capsid mutation N74D (HIV-1-N74D) or A77V (HIV-1-A77V), which fail to induce CPSF5 and CPSF6 translocation to nuclear speckles ^2,4^.

We employed a 3’-end sequencing approach, known as poly(A)-Click-sequencing (PAC-seq), to map global mRNA 3’-end changes in HIV-1–infected and mock-infected A549 cells. The PAC-seq technique is specifically designed to capture, enrich, and sequence polyadenylation tail junctions, enabling the direct identification and quantification of APA events rather than inferring their locations and assessing relative changes, which is the typical approach used when re-analyzing standard RNA-sequencing databases ^41,42^. We utilized our robust PolyAMiner analysis pipeline, which was custom-designed to analyze PAC-seq data by employing a vector projection–based engine to calculate intermediate polyadenylation signal usage ^43^ ^44^.

We first used differential gene expression analysis using DESeq2 to compare the PAC-seq results for RNA isolated from A549 cells infected with wild-type and mutant HIV-1. We found very similar results to those obtained using standard RNA-sequencing analyses (Figure S2A–C), showing significant upregulation of antiviral genes (Figure S2D) and significant downregulation of genes involved in chromosome maintenance and cell division (Figure S2E). We then analyzed PAC-seq data using PolyAMiner and found similar numbers of total mapped polyadenylation sites for each virus (Table S1). HIV-1 infection of A549 cells resulted in significant APA events, including both 3’UTR shortening and lengthening, compared with mock-infected controls (Figure 2A).

**Figure 2.**
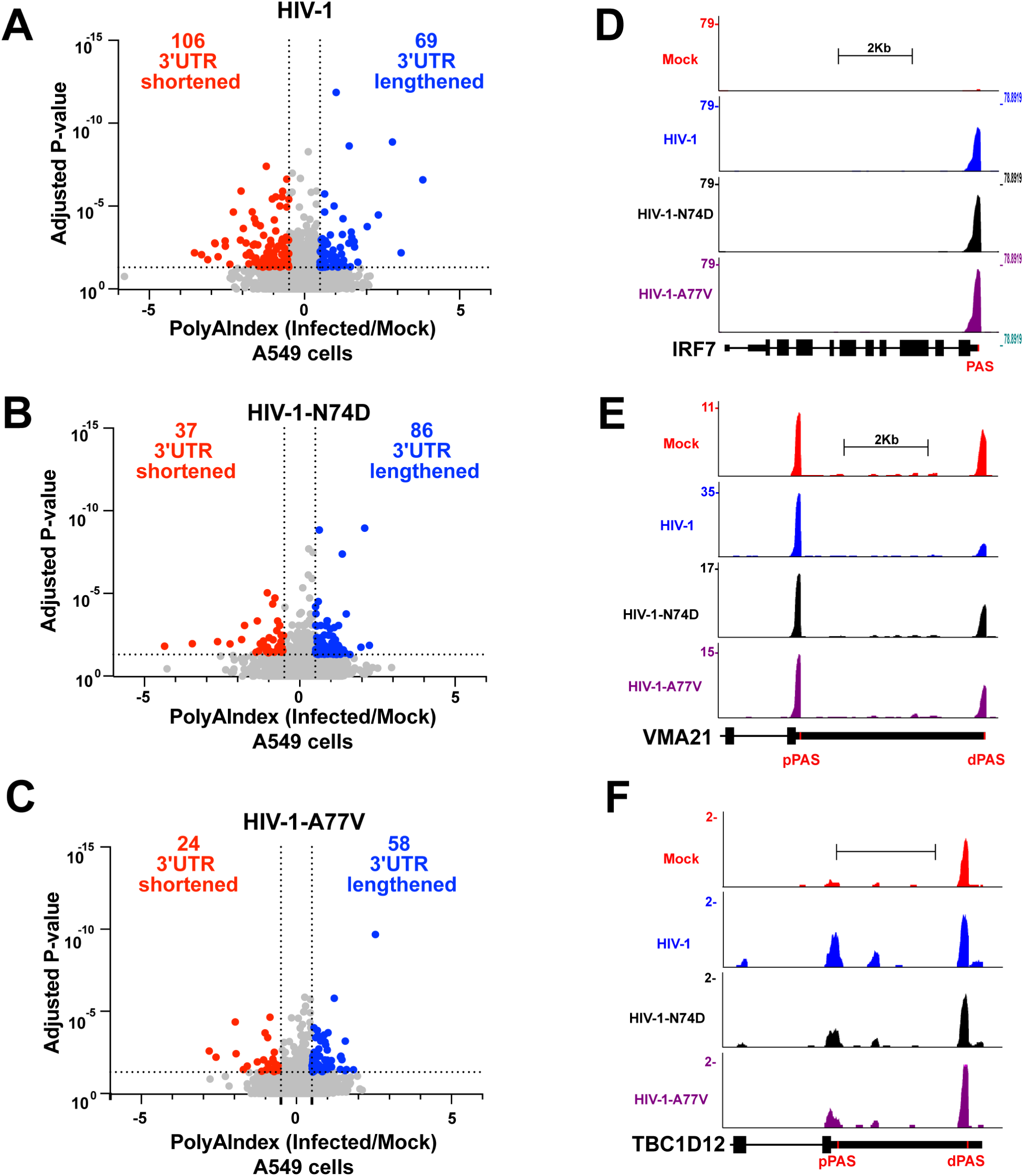
HIV-1 infection induces APA changes in human A549 cells that depend on the interaction between the viral capsid and CPSF6. (A-C) Human A549 cells were challenged with HIV-1-GFP **(A)**, HIV-1-N74D-GFP **(B)**, or HIV-1-A77V-GFP **(C),** using an MOI of 2 for 48 h. Cells were harvested, and total RNA was extracted. Total RNA from three infected and three mock-infected samples were sequenced to analyze changes in APA using PAC-seq. Scatterplots were generated from PolyAMiner analysis of PAC-seq results, plotted as the PolyAIndex (PAI) of HIV-infected cells relative to mock-infected cells. Negative PAI denotes 3’UTR shortening, whereas positive PAI denotes 3’UTR lengthening. Significance is defined as PAI<−0.5 or PAI>0.5, with an adjusted p-value<0.05. **(D–F)** Genome browser images of *IRF7* (D), *VMA21* (E), and *TBC1D12* (F) with PAC-seq reads shown. Scale is set to 2 Kb, and the y-axis shows read values. The position of the annotated polyadenylation signal (PAS) is shown in red. APA, alternative polyadenylation; GFP, green fluorescent protein; IRF7, interferon responsive factor 7; MOI, multiplicity of infection; PAC-seq, Poly(A)-Click-Sequencing; TBC1D12, TBC1 domain family member 12; UTR, untranslated region; VMA21, vacuolar membrane protein 21.

Compared with wild-type HIV-1 infections, infections with HIV-1-N74D or HIV-1-A77V, which do not induce CPSF5 and CPSF6 translocation to nuclear speckles, resulted in a similar number of 3’UTR lengthening events but a reduced number of 3’UTR shortening events (Figure 2B and C). These data suggest that HIV-1 infection partially mimics the CPSF5 KO phenotype, which results in a significant increase in the number of mRNA transcripts with shortened 3’UTRs ^36^. Functional gene analysis indicated an expected increase in antiviral gene expression, such as IFN regulatory factor 7 (Figure 2D), and shortened 3’UTRs for both vascular membrane protein 21 (VMA21, Figure 2E) and TBC1 domain family member 12 (TBC1D12, Figure 2F), although the shifts toward proximal polyadenylation sites for both VMA21 and TBC1D12 were less pronounced following infection with mutant HIV-1 than with wild-type HIV-1. These experiments indicate that HIV-1 infection–induced changes in APA on the capsid–CPSF6 interaction and that HIV-1 infection leverages the viral capsid–CPSF6 interaction to modulate the function of the CFIm complex.

### HIV-1 infection of human primary CD4^+^ T cells induces changes in alternative polyadenylation mediated by the interaction between the viral capsid and CPSF6

To investigate whether HIV-1 infection of human CD4^+^ T cells affects CFIm complex function, we measured changes in APA in human primary CD4^+^ T cells infected with wild-type HIV-1, HIV-1-N74D, or HIV-1-A77V (all expressing GFP as an infection reporter) at an MOI of 2. After 48 h, GFP-positive cells were sorted, and RNA was isolated and subjected to PAC-seq and PolyAMiner analyses. Similar to the response observed in A549 cells, human primary CD4^+^ T cells showed significant increases in the number of mRNA transcripts with altered polyadenylation site selection in HIV-1–infected cells than in mock-infected cells (Figure 3A and TableS2A). We also observed fewer mRNAs with shortened 3’UTRs in cells infected with HIV-1-N74D (Figure 3B and Table S2B) or HIV-1-A77V (Figure 3C and Table S2C). In accordance with our data from A549 cells, although of a more limited magnitude, these results show that 3’UTR shortening events induced by HIV-1 infection of primary CD4^+^ T cells correlates with the interaction between the viral capsid and CPSF6.

**Figure 3.**
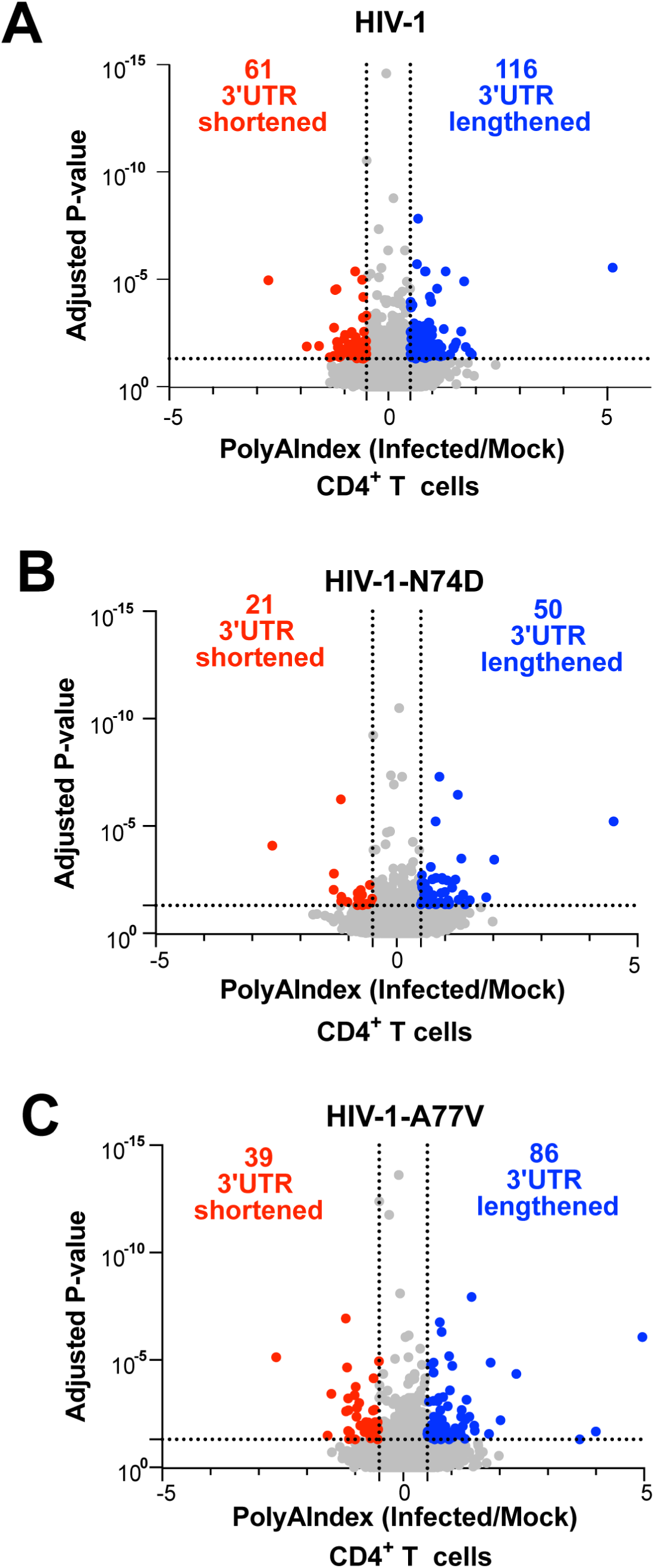
HIV-1 infection induces APA changes in human primary CD4^+^ T cells that depend on the interaction between the viral capsid and CPSF6. Human primary CD4^+^ T cells were challenged with HIV-1-GFP **(A)**, HIV-1-N74D-GFP **(B)**, or HIV-1-A77V-GFP **(C)** using an MOI of 2 for 48 h. After 48 h, cells were sorted to enrich for GFP-positive cells. Subsequently, GFP-positive cells were grown until a total of 96 h of HIV-1 infection was reached. Cells were harvested, and total RNA was extracted. Total RNA from three infected and three mock-infected samples was sequenced to analyze changes in APA using PAC-seq. Scatterplots were generated using PolyAMiner analysis of PAC-seq results, plotted as the PolyAIndex (PAI) of HIV-infected cells relative to mock-infected cells. A negative PAI denotes 3’UTR shortening in infected cells, whereas a positive PAI denotes 3’UTR lengthening. Significance was defined as PAI<−0.5 or PAI>0.5, with an adjusted p-value of <0.05. APA, alternative polyadenylation; GFP, green fluorescent protein; MOI, multiplicity of infection; PAC-seq, Poly(A)-Click-sequencing; UTR, untranslated region.

### HIV-1 infection–induced changes in alternative polyadenylation result in altered cellular protein expression

APA is a post-transcriptional regulatory mechanism that allows a single gene to produce mRNA transcripts with varying 3’UTR lengths. Even when the overall mRNA levels remain unchanged, the effects of different 3’UTR lengths become obvious at the protein level, as the 3’UTR contains regulatory elements, such as binding sites for RNA-binding proteins and microRNAs, that can significantly influence the gene expression pattern by altering mRNA stability, localization, or translational efficiency. Therefore, we sought to evaluate whether HIV-1 infection alters the expression of proteins encoded by mRNAs with varying 3’UTR lengths (Table 1), including those that displayed changes in transcript levels (Table 1).

**Table 1.** Top 10 Differentially Expressed Genes (DEGs) with altered 3’UTR lengths due to Alternative Polyadenylation (APA) in A549 cells infected with HIV-1 compared to mock-infected A549 cells, as determined by RNA-seq and REPAC analysis, as described in Methods.

| **SYMBOL** | **Gene ID** | **log2FoldChange** | **% Fold Change** | **Adjusted p value** | **PolyA** |
| --- | --- | --- | --- | --- | --- |
| **Downregulated** |  |  |  |  |  |
| **FOXP1** | 27086 | -2.4610 | 81.84 | 0.00284 | Short |
| **KIF20B** | 9585 | -1.3494 | 60.75 | 0.00087 | Short |
| **LDLRAD4** | 753 | -1.2530 | 58.04 | 0.03106 | Short |
| **STRIP2** | 57464 | -1.1298 | 54.30 | 0.00183 | Long |
| **WDHD1** | 11169 | -1.0417 | 51.43 | 4.90875e-05 | Short |
| **Upregulated** |  |  |  |  |  |
| **TP53INP1** | 94241 | 1.8110 | 250.89 | 6.33916e-07 | Short |
| **SLFN5** | 162394 | 1.7320 | 232.18 | 1.52665e-06 | Short |
| **MSMO1** | 6307 | 1.3008 | 146.37 | 2.16076e-05 | Short |
| **MR1** | 3140 | 1.2569 | 138.98 | 0.00013 | Short |
| **MDM2** | 4193 | 1.0620 | 108.79 | 1.26122e-08 | Long |

Our analysis revealed that the transcript encoding Schlafen family member 5 (SLFN5) was upregulated upon HIV-1 infection but with a shortened 3’UTR (Table 1) due to changes in APA upon HIV-1 infection (Figure 4A). Therefore, SLFN5 was selected as a promising candidate to explore the effects of changes in 3’UTR length on protein expression. We infected human A549 cells with HIV-1-GFP at an MOI of 2 for 48 h and assessed protein expression by western blot analysis. Across three independent HIV-1-GFP preparations, HIV-1 infection increased SLFN5 protein expression by 6–7 fold that of mock-infected cells (Figure 4B). We also infected A549 cells with an HIV-1 expressing luciferase (HIV-1-Luc) at an MOI of 2 for 48 h, and immunoblot analysis revealed that infected cells had significantly increased SLFN5 levels that were 7–8-fold the levels in mock-infected controls (Figure 4C). We also measured SLFN5 expression upon infection with HIV-1-N74D or HIV-1-A77V, which express a capsid protein that does not interact with CPSF6, and found that these mutant viruses failed to induce SLFN5 expression (Figure 4D), suggesting that this induction is dependent on the interaction between the capsid protein and CPSF6. To verify that the observed increase in SLFN5 expression was specifically due to HIV-1 infection, we repeated the experiment using heat-inactivated HIV-1-GFP (95°C for 30 min). We found that the SLFN5 levels in cells exposed to heat-inactivated virus were comparable to the levels in mock-infected cells (Figure 4E), confirming that active HIV-1 infection is necessary to induce SLFN5 expression.

**Figure 4.**
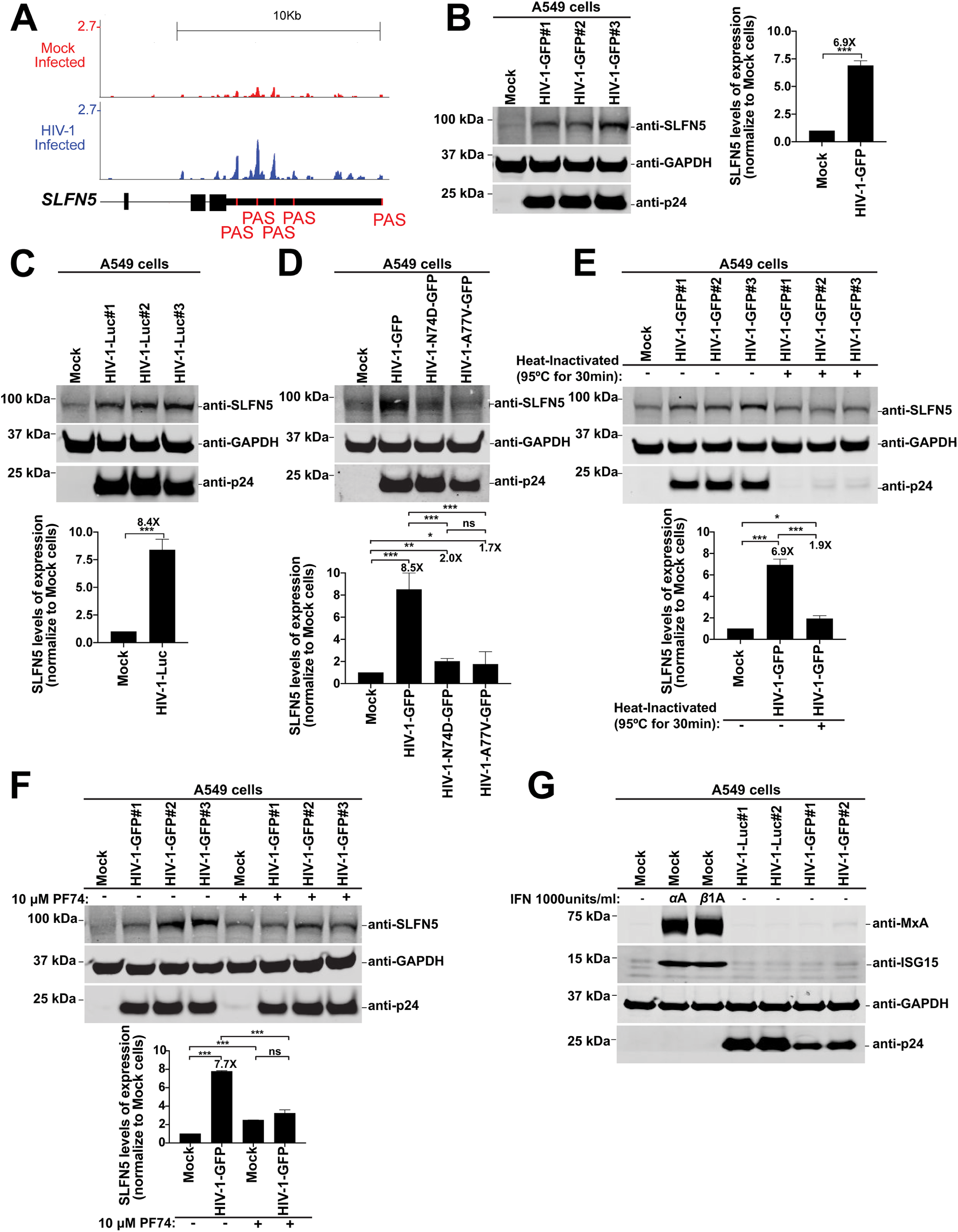
HIV-1 infection induces changes in protein expression levels. **(A)** Genome browser images of *SLFN5* in HIV-infected or mock-infected A549 cells with PAC-seq reads shown. Scale is set to 10 Kb, and the y-axis shows read values. The position of the annotated polyA signal is shown in red. **(B–G)** Human A549 cells were infected under the following conditions: **(B)** three independent HIV-1-GFP preparations at an MOI of 2 for 48 h; **(C)** three independent HIV-1-Luc preparations at an MOI of 2 for 48 h; **(D)** HIV-1-GFP, HIV-1-N74D-GFP, or HIV-1-A77V-GFP at an MOI of 2 for 48 h; **(E)** HIV-1-GFP with or without heat-inactivation (95°C for 30 min) at an MOI of 2 for 48 h; **(F)** HIV-1-GFP at an MOI of 2 for 48 h with or without 10 µM PF74; and **(G)** HIV-1-Luc or HIV-1-GFP at an MOI of 2 for 48 h. Cells were lysed, and proteins were analyzed by western blot using the indicated antibodies. Virus presence was assessed using anti-p24 antibodies, and anti-GAPDH antibodies were used as a protein loading control. Each experiment was repeated at least three times, and a representative image is shown. Graphs show the average densitometry quantification of three replicates with standard deviation. Significance was determined using unpaired t-test; *p<0.05; **p<0.01; ***p<0.001; ns, not significant. GAPDH, glyceraldehyde 3-phosphate dehydrogenase; GFP, green fluorescent protein; ISG15, interferon-stimulated gene 15; Luc, luciferase; MOI, multiplicity of infection; MxA, human myxovirus resistance protein 1; p24, viral capsid; SLFN5, Schlafen family member 5.

To determine whether the increased SLFN5 expression observed in HIV-1– infected cells correlates with CPSF5 and CPSF6 translocation to nuclear speckles, we analyzed SLFN5 expression in human cells infected with HIV-1 in the presence of the small molecule PF74, which prevents CPSF5 and CPSF6 translocation to nuclear speckles during infection ^45^. Similar to cells infected with HIV-1-N74D and HIV-1-A77V, cells infected with HIV-1 in the presence of PF74 failed to induce SLFN5 expression above the levels observed in mock-infected cells (Figure 4F). These findings suggest that the induction of SLFN5 expression in HIV-1–infected cells depends on the translocation of CPSF5 and CPSF6 to nuclear speckles, highlighting a potential link between capsid-dependent nuclear events and SLFN5 expression.

Because SLFN5 is a type I IFN-induced protein, we tested whether IFN-inducing contaminants in the viral preparation could explain the observed SLFN5 upregulation. A549 cells were infected with HIV-1-GFP at an MOI of 2 for 48 h, and the protein levels of myxovirus resistance protein 1 (MxA) and IFN-stimulated gene 15 (ISG15), which are both type I IFN-induced proteins, were analyzed by western blot. As a positive control, uninfected cells were treated with 1000 U/mL of IFNα or IFNβ for ∼40 h. The protein expression levels of MxA and ISG15 in infected cells were comparable to the levels observed in mock-infected controls (Figure 4G), indicating the absence of IFN-inducing contaminants. These findings demonstrate that HIV-1 infection itself drives the upregulation of SLFN5. Overall, these experiments indicate that the protein expression levels of SLFN5 are upregulated as a result of HIV-1–induced changes in APA.

### The loss of CPSF6 expression shortens the 3’UTRs of cellular mRNAs at a global level

Our results suggest that HIV-1 infection affects APA through the interaction of viral capsid with CPSF6, resulting in the shortening of mRNA 3’UTRs, ultimately changing cellular protein expression levels. The absence of CPSF5 or CPSF6 expression in different cell types results in the global shortening of all cellular mRNA 3’UTRs ^11,14,31,32,36–38^; therefore, we decided to test whether knocking out CPSF6 in A549 cells would also increase the number of cellular mRNAs with shorter 3’UTRs. We used the CRISPR-Cas9 system to generate CPSF6-KO cells and control A549 cells using a non-targeting clone (NT#H1). From 12 single CPSF6-KO clones, we selected three independent clones for this study (Figure 5A), designated as CPSF6-KO#B4, CPSF6-KO#B7, and CPSF6-KO#C8, and confirmed total CPSF6 depletion in each clone by western blot. As controls, we used parental A549 cells (wild-type) and the NT#H1 cells. We also evaluated the expression levels of other CFIm complex members, such as CPSF5 and CPSF7. We found a significant decrease in CPSF5 expression levels in CPSF6-KO clones compared with the levels in control cells (Figure 5A). By contrast, a modest increase in CPSF7 expression levels was observed in the CPSF6-KO cell lines compared with the levels in wild-type cells (Figure 5A). These experiments suggest the possibility that CPSF5 stability depends on CPSF6 expression, either because CPSF5 and CPSF6 form a heterotetramer or because any impact on the CFIm complex affects CPSF5 expression. Together, these results suggest that a regulatory mechanism may exist among the expression levels of different CFIm complex members.

**Figure 5.**
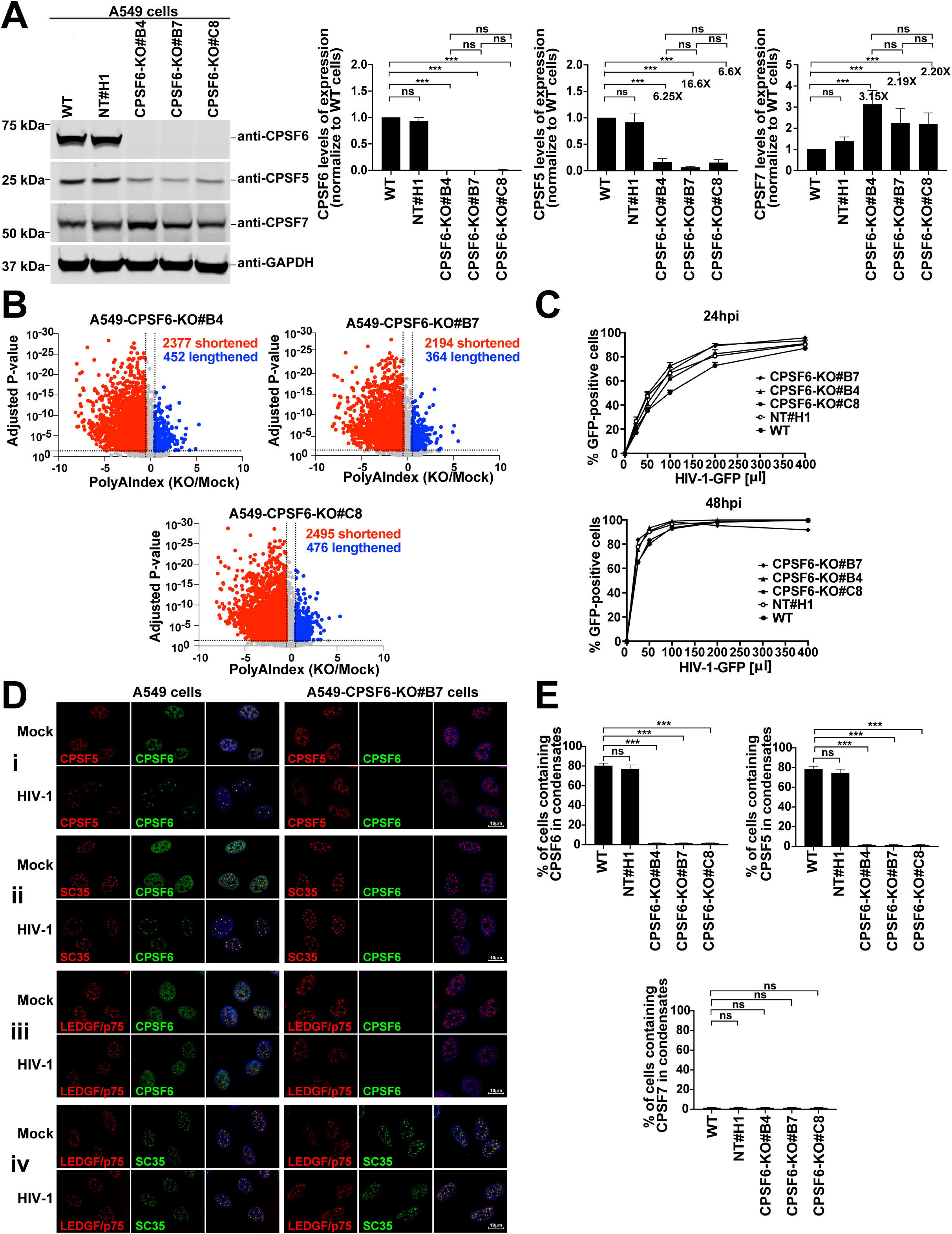
Loss of CPSF6 expression globally shortens the 3’UTR of cellular mRNAs. **(A)** A549 WT, NT#H1, CPSF6-KO#B4, CPSF6-KO#B7, and CPSF6-KO#C8 cells were lysed, and proteins were analyzed by Nu-PAGE, followed by western blot using anti-CPSF6, anti-CPSF5, anti-CPSF7, and anti-GAPDH antibodies. Experiments were repeated at least three times, and a representative image is shown. Graphs show the average densitometry quantification of three replicates with standard deviation. Significance was determined using unpaired t-test; *p<0.05; **p<0.01; ***p<0.001; ns, not significant. **(B)** Total RNA from CPSF6-KO and WT A549 cells was prepared, and polyadenylated transcripts were identified by PAC-seq, followed by APA analysis using PolyAMiner. **(C)** A549 WT, NT#H1, CPSF6-KO#B4, CPSF6-KO#B7, and CPSF6-KO#C8 cells were infected with increasing amounts of HIV-1-GFP for 24 or 48 h. Infection was assessed as the percentage of GFP-positive cells by flow cytometry. **(D)** WT and CPSF6-KO A549 cells were infected with HIV-1-GFP at an MOI of 2 for 48 h. Cells were fixed, permeabilized, and stained using the following antibodies: (i) anti-CPSF5 (red) and anti-CPSF6 (green); (ii) anti-SC-35 (red) and anti-CPSF6 (green); (iii) anti-LEDGF/p75 (red) and anti-CPSF6 (green); and (iv) anti-LEDGF/p75 (red) and anti-SC35 (green). Nuclei were stained with DAPI (blue). Scale bar, 10 µm. **(E)** Percentage of A549 cells containing CPSF6, CPSF5, or CPSF7 in nuclear speckles (condensates) upon HIV-1 infection (average of three independent experiments with standard deviation). Cells containing CPSF6, CPSF5, or CPSF7 in nuclear speckles were determined by visual examination of 200 cells APA, alternative polyadenylation; CPSF5, cleavage and polyadenylation specificity factor subset 5; CPSF6, cleavage and polyadenylation specificity factor subset 6; CPSF7, cleavage and polyadenylation specificity factor subset 7; DAPI, 4’,6-diamidino-2-phenylindole; GAPDH, glyceraldehyde 3-phosphate dehydrogenase; KO, knockout; LEDGF/p75, lens epithelium-derived growth factor; MOI, multiplicity of infection; NT, non-targeting; PAC-Seq, Poly(A)-Click-Sequencing; UTR, untranslated region; WT, wild-type; hpi, hours post-infection.

We also used PAC-seq to analyze changes in APA in CPSF6-KO cells (Figure 5B and Table S3A-C). Consistent with the known role of CPSF5 in APA, we observed a global shortening of mRNA 3’UTRs in CPSF6-KO cells compared with those in wild-type cells. Because CPSF5 and CPSF6 work together in the CFIm complex, these findings align with previous reports that the reduction or complete depletion of CPSF5 in human cells increases the prevalence of mRNAs with shorter 3’UTRs ^23,36,46^. Consistently, we observed that the expression of VMA21, a well-established reporter of changes in alternative polyadenylation (APA) ^47^, was increased in CPSF6-KO cells compared to wild-type cells, suggesting that CPSF6-KO leads to a complete deregulation of cellular APA (Figure S3).

Next, we tested the effects of CPSF6 depletion on HIV-1 infectivity in A549 cells. We challenged CPSF6-KO and control cells with increasing amounts of HIV-1-GFP for 24 or 48 h and determined infectivity by measuring the percentage of GFP-positive cells using flow cytometry (Figure 5C). These results showed that CPSF6 depletion does not significantly affect HIV-1 infection in A549 cells, similar to previous reports in human primary cells and other cell lines ^10,13,14^. Previously, we and others described that HIV-1 infection induces the translocation of both CPSF6 and CPSF5, a CFIm complex member that works together with CPSF6, to nuclear speckles ^5^. Therefore, we next examined whether HIV-1 infection can induce CPSF5 translocation to nuclear speckles in CPSF6-KO cells. CPSF6-KO A549 cells were infected with HIV-1-GFP at an MOI of 2 for 48 h, fixed, permeabilized, and subjected to immunofluorescence analysis using anti-SC35, anti-CPSF5, and anti–lens epithelium-derived growth factor/p75 (LEDGF/p75) antibodies. Unlike in wild-type cells, HIV-1 infection did not induce CPSF5 translocation to nuclear speckles in CPSF6-KO cells, suggesting that CPSF5 translocation to nuclear speckles is dependent on CPSF6 translocation (Figure 5Di and 5E). However, the localization of the nuclear speckle marker SC35 was unchanged between wild-type and CPSF6-KO A549 cells, indicating that nuclear speckles were not affected (Figure 5Dii and 5E). We previously showed that the HIV-1 integration cofactor LEDGF/p75 surrounds nuclear speckles containing CPSF5 and CPSF6 upon HIV-1 infection ^5^; therefore, we tested whether CPSF6 was required for this process to occur and found that LEDGF/p75 does not surround nuclear speckles upon HIV-1 infection in CPSF6-KO cells (Figure 5Diii and 5Div). As a control, we quantified the percentage of cells containing CPSF5, CPSF6, and CPSF7 in nuclear speckles upon HIV-1 infection of wild-type and CPSF6-KO cells (Figure 5E). Overall, these results indicate that the translocation of CPSF6 to nuclear speckles is important for the recruitment of CPSF5 and LEDGF/p75 to nuclear speckles upon HIV-1 infection.

### HIV-1 infection mimics the loss of CPSF6 expression

The analysis of protein expression levels encoded by mRNA transcripts undergoing APA revealed that SLFN5 protein levels are upregulated upon HIV-1 infection. Because HIV-1 infection mimics the phenotype observed in CPSF6-KO cells, we investigated whether CPSF6 depletion influences SLFN5 expression. CPSF6-KO cells exhibited high SFLN5 levels, up to 20-fold the levels observed in wild-type or NT#H1 control cells (Figure 6A), similar to the upregulation observed during HIV-1 infection. These experiments suggest that SLFN5 expression levels are regulated by APA.

**Figure 6.**
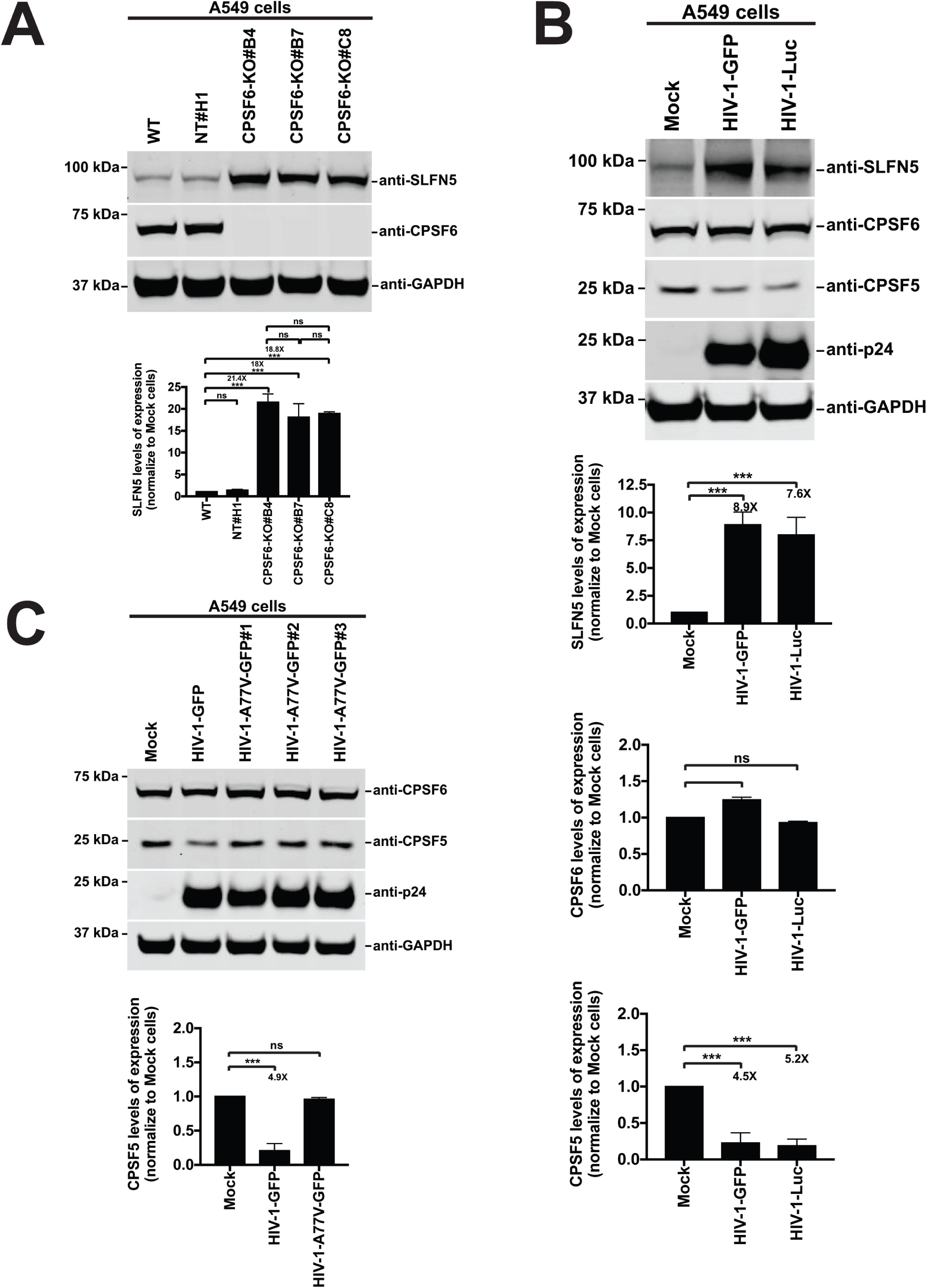
HIV-1 infection mimics the CPSF6-KO phenotype. **(A)** CPSF6-KO and control (WT and NT#H1) A549 cells were analyzed by western blot using anti-SLFN5 and anti-CPSF6 antibodies. Anti-GAPDH antibodies were used as a loading control. **(B)** A549 cells were challenged with HIV-1-GFP or HIV-1-Luc using an MOI of 2 for 48 hours. Subsequently, cells were lysed, and extracts were analyzed by western blot using anti-SLFN5, anti-CPSF6, anti-CPSF5, anti-p24, and anti-GAPDH antibodies. **(C)** A549 cells were challenged with three different HIV-1-A77V-GFP preparations using an MOI of 2 for 48 hours. Cells were lysed and analyzed by western blot using anti-SLFN5, anti-CPSF5, and anti-p24. GAPDH was used as a loading control. (**A–C**) All experiments were repeated at least three times, and a representative image is shown. Graphs show the average densitometry quantification of at least three replicates with standard deviation. Significance was determined using unpaired t-test; *p<0.05; **p<0.01; ***p<0.001; ns, not significant. CPSF5, cleavage and polyadenylation specificity factor subset 5; CPSF6, cleavage and polyadenylation specificity factor subset 6; GAPDH, glyceraldehyde 3-phosphate dehydrogenase; GFP, green fluorescent protein KO, knockout; Luc, luciferase; MOI, multiplicity of infection; NT, non- targeting; p24, viral capsid; SLFN5, Schlafen family member 5; WT, wild-type.

Because we found that CPSF6-KO cells exhibit a consistent decrease in CPSF5 protein expression compared with wild-type cells, we tested whether HIV-1 infection decreases CPSF5 expression. Similar to the phenotype observed in CPSF6-KO cells, we observed that HIV-1 infection decreases CPSF5 protein expression levels (Figure 6B). To test the specificity of HIV-1 infection–induced decreases in CPSF5 protein expression, we measured CPSF5 levels upon infection with HIV-1-A77V, which does not induce the translocation of CPSF5 and CPSF6 to nuclear speckles. Unlike the decrease in SLFN5 levels observed with wild-type HIV-1 infection, CPSF5 expression levels in cells infected with HIV-1-A77V were similar to those in mock-infected cells (Figure 6C), suggesting that HIV-1 infection specifically alters cellular expression patterns through the interaction of the viral capsid with CPSF6.

## DISCUSSION

We and others have shown that HIV-1 infection triggers the translocation of CPSF6 to nuclear speckles ^2,4^. We also demonstrated that the HIV-1 core must enter the nuclear compartment for this translocation to occur and that this process is dependent on the viral capsid protein. Interestingly, CPSF5 also translocates to nuclear speckles upon HIV-1 infection ^5,14,48^. Over the years, multiple groups have demonstrated that CPSF6 depletion does not significantly impair HIV-1 infectivity, suggesting that CPSF6 may not be essential for productive infection, likely because the virus utilizes alternative pathways in the absence of CPSF6, leaving the precise role played by CPSF6 during HIV-1 infection unclear. We recently speculated that HIV-1 induces the translocation of CPSF6 to nuclear speckles as a strategy for regulating cellular gene expression ^35^. Because both CPSF5 and CPSF6 are involved in the regulation of APA, a key cellular mechanism that controls gene expression, we tested the hypothesis that HIV-1 infection alters the nuclear localization of CPSF5 and CPSF6, thereby dysregulating APA and ultimately modifying cellular gene expression.

To understand the ability of HIV-1 to modulate cellular gene expression in human cells, we first used RNA-sequencing to define all gene expression changes triggered by HIV-1 infection in human primary cells and cell lines. Using different databases, we identified the activation of genes countering viral infection (response to virus, viral processes, and others) and the suppression of genes involved in DNA metabolism (DNA replication, recombination and repair, and cell division). HIV-1 infection of A549 cells led to more genes being significantly downregulated than upregulated, which is consistent with previous RNA-sequencing studies of HIV-1–infected cells ^49^.

To test the hypothesis that HIV-1 may impact gene expression via APA modulation, we measured APA in HIV-1–infected human A549 cells and human primary CD4^+^ T cells using two independent, state-of-the-art methodologies: PAC-seq, which maps global mRNA 3’-end changes by specifically sequencing polyadenylated transcripts ^41^, followed by PolyAMiner ^43^; and REPAC ^39^, an algorithm that uses total RNA-sequencing data to identify changes in 3’UTRs. Our analyses revealed that HIV-1 infection globally modulates APA, leading to a significant increase not only of mRNAs with shorter 3’UTRs but also, to a lesser extent, longer 3’UTRs. The mRNA 3’UTR contains regulatory elements that determine mRNA metabolic features, including stability, translation, nuclear export, and cellular localization ^25^. Therefore, changing the 3’UTR length through the selection of APA sites can dramatically affect gene expression and function. These results demonstrate that HIV-1 uses APA as at least one mechanism for controlling gene expression. We also showed that infection by HIV-1 bearing a capsid mutation that prevents the interaction of the viral capsid with CPSF6 was unable to modulate APA, suggesting that the ability of the viral capsid to interact with CPSF6 is essential for modulating cellular APA.

This study suggests that HIV-1 infection–induced translocation to nuclear speckles affects the cellular functions of CPSF5 and CPSF6. In support of this finding, CPSF5 depletion in human cells alters APA regulation ^36,46^, ultimately impacting cellular gene expression. Thus, the interaction of the viral core with CPSF6 within the nuclear compartment likely represents an upstream event that leads to changes in cellular gene expression. In agreement with these results, similar to CPSF5 depletion, CPSF6 KO in human A549 cells significantly increases the number of mRNAs with short 3’UTRs. The ability of viruses to modulate host gene expression to facilitate viral replication is well established, with numerous viruses from various families known to manipulate cellular gene expression through viral proteins that target transcription, mRNA processing, and translation. For example, HSV-1 inhibits splicing via the viral protein ICP27, poliovirus blocks mRNA export via the protein 2A, influenza virus suppresses polyadenylation via NS1, adenovirus inhibits transcription initiation via E1A, rubella virus impedes translation via its capsid protein, and Bunya viruses inhibit translation via their NS proteins. These examples illustrate that viruses have evolved diverse strategies for controlling the gene expression profiles of infected cells. In most cases, the viral regulation of host gene expression ultimately serves to create a permissive environment for viral replication. We propose that HIV-1 infection alters cellular expression by modulating APA through the interaction of the viral capsid with CPSF6.

Depleting CPSF5 expression in human cells significantly increases the number of cellular mRNAs with shorter 3’ UTRs ^36,46^. To determine whether CPSF6 depletion has similar effects, we generated CPSF6-KO A549 cells and analyzed changes in APA. Consistent with the results of CPSF5 depletion ^36^, CPSF6-KO cells displayed a marked increase in mRNAs with shorter 3’ UTRs. CPSF5 and CPSF6 function together within the CFIm complex, and the loss of either protein is likely to disrupt the integrity of this complex, severely impacting APA. Our observations also revealed that CPSF5 and LEDGF/p75 translocation to nuclear speckles is dependent on CPSF6 expression, suggesting that the interaction of the viral capsid with CPSF6 occurs upstream from the recruitment of CPSF5 and LEDGF/p75 to nuclear speckles. HIV-1 infection modulates the CFIm complex through direct interaction between the viral capsid and CPSF6, and these experiments suggest that the sequestration of CPSF6 to nuclear speckles by the viral capsid affects the cellular function of the CFIm complex.

To identify a protein that could potentially be used to monitor changes in APA during HIV-1 infection, we examined several proteins encoded by transcripts that undergo APA in HIV-1–infected cells. SLFN5 expression increases during HIV-1 infection and in CPSF6-KO cells due to APA changes. These findings suggest that SLFN5 may serve as a useful tool for monitoring HIV-1–induced changes in APA. We observed that both HIV-1 infection and CPSF6-KO reduce CPSF5 expression levels in human cells. Overall, this study suggests that HIV-1 infection mimics the phenotype observed in CPSF6-KO cells, indicating that HIV-1 sequesters CPSF5 and CPSF6 in nuclear speckles to alter their function, representing a novel regulatory mechanism.

## MATERIALS AND METHODS

### Cell culture and generation of cell lines

Human lung carcinoma A549 and human embryonic kidney HEK293T cells were obtained from the American Type Culture Collection (A549: Cat # CCL-1885; HEK293T: Cat #CRL-3216) and maintained in Dulbecco’s modified Eagle medium supplemented with 10% heat-inactivated fetal bovine serum (FBS), 100 U/ml penicillin, 100 µg/ml streptomycin, 29.2 mg/ml L-glutamine (Life Sciences), and 5 µg/ml plasmocin (Invivo Gen, San Diego, CA), in a humidified incubator with 5% CO_2_ at 37°C. Human lymphocytes were isolated from peripheral blood mononuclear cells obtained from three healthy donors using the Pan T Cells Isolation kit (Miltenyi Biotec, Cat#130-096-535). T cells were activated by culturing in complete Roswell Park Memorial Institute (RPMI-1640) medium supplemented with 4 µg/ml phytohemagglutinin-L (PHA-L) and 20 IU/mL interleukin-2 (IL-2; Miltenyi). CPSF6-KO A549 cells were generated using the CRISPR-Cas9 ribonucleoprotein (crRNP) gene-editing system, according to the manufacturer’s instructions (Synthego). The sequences of the synthetic guide RNAs (sgRNAs) targeting CPSF6 were: sgRNA-1: 5’-CUUUUUAGGUUUGCCCUUGU −3’, sgRNA-2: 5’-UUACCUAAAAGAGAACUUCA-3’, and sgRNA-3: 5’-UGAGGACUGCUUACUUUUCC-3’. Cells were electroporated with Amaxa 4D nucleofector according to the manufacturer’s instructions (Lonza). After 72 h, cells were subjected to limiting dilution in 96-well plates. Clones were assessed for KO by immunofluorescence and immunoblot analysis with rabbit polyclonal CPSF6 antibody. Control cells were nucleofected with crRNPs carrying a non-targeting sgRNA (NT#H1): 5’-GCACUACCAGAGCUAACUCA-3’.

### Antibodies and cell reagents

We used mouse monoclonal antibodies targeting the following proteins: SC35 (clone SC-35; Cat# ab11826, Abcam), CPSF5 (clone 3F8;Cat# H00011051-M12, Novus Biologicals), CPSF7 (clone A-9; Cat# sc-393880, Santa Cruz), CPSF6 (clone F-3; Cat# sc-376228, Santa Cruz), ISG15 (clone F-9; Cat# sc-166755, Santa Cruz), and HIV-1 p24 (clone 183-H12-5C; Cat# ARP-3537, NIH AIDS Reagent Program). We used rabbit polyclonal antibodies against the following proteins: CPSF6 (Cat# ab99347, Abcam), LEDGF/p75 (Cat# A300-847A, Bethyl Laboratories, Inc), SLFN5 (Cat#PA5-53638, Invitrogen), and MxA (Cat#PA5-22101, Invitrogen) and VMA21 (Cat#21921-1-AP, Proteintech). The fluorescent nuclear stain 4’,6-diamidino-2-phenylindole (DAPI), and the following fluorescently labeled antibodies were from Life Technologies: Alexa Fluor 488-conjugated, Alexa Fluor 594-conjugated, and Alexa Fluor 647-conjugated donkey anti-rabbit IgG and anti-mouse IgG. The human IFNαA and 1βA (Cat#IF007 and IF014, respectively) were from Millipore. TRIzol® reagent (Cat#15596026) was from Invitrogen. Dimethyl sulfoxide (DMSO) (Cat# D2438), PF74 (Cat#SML0835), and chloroform (Cat#C298-1) were from Sigma-Aldrich.

### Production of HIV-1

Wild-type and mutant HIV-1 (HIV-1-N74D and HIV-1-A77V) expressing GFP or Luc as reporters were produced by co-transfecting HIV-1-gag-pol, LTR-GFP-LTR/LTR-Luc-LTR, tat, rev, and VSV-G in HEK293T/17 cells, as described ^50^. Viruses were collected 48 h after transfection, filtered, concentrated, tittered, aliquoted, and stored at −80°C. Heat-inactivation of HIV-1 was performed by incubating the viral preparations at 95°C for 30 min.

### Analysis of viral infectivity by flow cytometry

The infectivity and titer of HIV-1-Luc were measured using TZM-bl GFP-reporter cells, in which HIV-1 infection induces GFP, as described ^51^. We infected human A549 cells and determined the percentage of GFP-positive cells using a flow cytometer (BD Celesta).

Virus titer calculation was performed according to the following equation: Infectious units (IU)/ml = (cell number) × (% of GFP-positive cells) × (dilution factor), where the dilution factor = 1000 µl ÷ viral input (µl). To calculate the volume of virus used at a specific MOI, use the following equation: MOI = [(virus stock IU/ml) × (volume of virus used)] ÷ (number of cells in infection).

### Immunofluorescence microscopy image acquisition and deconvolution

Samples for immunofluorescence analysis were prepared as described previously, with some modifications ^2^. Briefly, cells were seeded on 12-mm, round, glass coverslips in a 24-well plate and maintained in a complete culture medium. After HIV-1 infection, the coverslips were rinsed with phosphate-buffered saline (PBS) and fixed with 4% paraformaldehyde-PBS for 15−30 min at room temperature. Subsequently, cells were incubated in 0.1 M glycine-PBS for 10 min at room temperature. Cells were permeabilized using 0.5% Triton X-100 for 5 min at room temperature. Non-specific binding was prevented using a blocking solution (3% bovine serum albumin in PBS) for 60 min at room temperature. Samples were incubated with primary antibodies in a blocking solution for 60 min in a dark room at room temperature. Subsequently, the coverslips were rinsed with PBS and incubated with the appropriate secondary antibodies against mouse or rabbit IgG in a blocking solution for 30 min at room temperature. Finally, the coverslips were washed with PBS and mounted using FluorSave reagent (Sigma-Aldrich). Fluorescence microscopy images were acquired with an AxioObserver.Z1 microscope equipped with a PlanApo 63× oil immersion objective (NA 1.4) and an AxioCam MRm digital camera (Carl Zeiss). Image acquisition was performed using a Zeiss Z1 Observer inverted microscope and ZEN 3.3 (blue edition) software. Image deconvolution was performed with the ZEN 3.3 software using an acquired point spread function. Images for figures were processed with Adobe Photoshop CS5 software (Adobe Systems, Mountain View, CA).

### Protein extracts and immunoblotting

Preparation of protein extracts from cultured cells was performed using a method that we have described elsewhere ^2^. Briefly, cells were washed twice with ice-cold PBS and incubated at 4°C for 1 h in whole-cell extract buffer (50 mM Tris-HCl, 280 mM NaCl, 0.5% IGEPAL, 0.2 nM EDTA, 2 mM EGTA, 10% glycerol, and 1 mM DTT, pH 8) supplemented with a cocktail of protease inhibitors, benzonase, and 50 µg/ml ethidium bromide. Lysates were cleared by centrifugation at 14,000×*g* for 60 min at 4°C. Protein concentration was determined with a protein assay dye reagent (Bio-Rad Laboratories, Hercules, CA, USA). Depending on the protein analyzed, samples of protein extracts were denatured at 37°C or 65°C for 10 or 60 min, respectively, in 4× Nu-PAGE LDS sample buffer (Cat#NP0007, Invitrogen) and separated by Nu-PAGE. After electrophoresis, proteins were transferred to a nitrocellulose membrane and incubated sequentially with corresponding primary and secondary antibodies overnight at 4°C and 1 h at room temperature, respectively. Detection of proteins was performed using the LI-COR Odyssey Imaging System (LI-COR, Biotech). As an internal gel loading control, we analyzed the levels of glyceraldehyde 3-phosphate dehydrogenase (GAPDH). The immunoblot signal from images with non-saturated pixels was estimated using ImageJ software (version 1.47h, Bethesda, MD). For each condition, protein bands were quantified from three independent experiments.

### RNA extraction

Total RNA from A549 mock-infected and HIV-1-infected cells (in triplicate) was extracted with TRIzol reagent (Invitrogen, CA, USA) according to the manufacturer’s protocol. Briefly, TRIzol Reagent was directly added to the culture dish to lyse the cells (∼5×10^6^ cells). The homogenized content was transferred to a 1.5-mL microtube and incubated for 5 min at room temperature. Chloroform was added to the sample, and the sample was shaken for 30 s and incubated for 3 min at room temperature, followed by centrifugation at 12,000×*g* for 15 min at 4°C. The aqueous phase containing the RNA was transferred to a new 1.5-mL microtube. For RNA precipitation, an equal volume of isopropanol (Sigma-Aldrich, St. Louis-MO–USA) was added to the sample, which was incubated at 4°C for 10 min. Next, the sample was centrifuged at 12,000×*g* for 10 min at 4°C, and the supernatant was discarded. The RNA pellet was washed using 75% (v/v) ethanol (Sigma-Aldrich, St. Louis-MO–USA), followed by centrifugation at 10,000×*g* for 5 min at 4°C. The supernatant was discarded, and the pellet was dried at room temperature. RNA was solubilized in RNase-free ultrapure water. The A230/260 and A260/280 ratios were measured by a NanoDrop spectrophotometer to determine the purity of the isolated total RNA. Isolated RNA was stored at −80°C until analysis.

### RNA-sequencing data processing

The RNA-sequencing data processing pipeline was adapted from published protocols. Briefly, we first trimmed the paired-end reads in FASTQ files using Trimmomatic (version 0.39)2, targeting known adapters (TruSeq3-SE.fa:2:30:10). We also trimmed the first three and last three base pairs if their quality fell below set thresholds.

Additionally, we employed a 4-base sliding window technique, trimming reads when the average window quality dropped below 15. Reads shorter than 36 base pairs were excluded. Next, the trimmed reads were aligned to the human reference genome GRCh38 (https://www.ncbi.nlm.nih.gov/datasets/genome/GCF_000001405.26/) using HISAT2 (PMID: 31375807, v 2.1.0) and converted into BAM format using samtools (version 1.6) ^52^. The alignment data was then processed by StringTie (version 2.1.3)4 to estimate gene expression levels. The StringTie merge function merged samples’ gene structures to create a unified set of transcripts across all samples. Finally, read counts were obtained using prepDE.py (https://ccb.jhu.edu/software/stringtie/index.shtml?t=manual) to prepare the data for DESeq2.

#### Differential gene expression analysis

The raw read counts were processed in R (version 4.2.3) using DESeq2 (version 1.38.3)5 to investigate differences across conditions, accounting for donor variability and batch effects. Genes with a minimum count of 10 in at least three samples were retained for analysis. To visualize treatment effects, counts were transformed using variance stabilizing transformation function from DESeq2 and subjected to principal components analysis, visualizing the first two principal components6. Differential gene expression between infected and mock-infected cells was assessed using the DESeq function from DESeq2, which estimates size factors and dispersions, fitting a negative binomial generalized linear model. P-values were adjusted using the Benjamini-Hochberg method (https://www.jstor.org/stable/2346101). Adjusted p-values below 0.05 were considered significant. Gene enrichment analysis was subsequently performed using gene set enrichment analysis (GSEA) to identify specific Gene Ontology, Kyoto Encyclopedia of Genes and Genomes, and Reactome pathways associated with HIV infection. We performed GSEA using clusterProfiler v.4.10.0 in R, with all gene lists ranked by the corresponding log2 fold-change. For these analyses, all genes whose gene symbols could be mapped to ENTREZ IDs using the org.Hs.eg.db v.3.18.0 Bioconductor annotation package were included.

#### APA Analysis using REPAC

APA analysis was conducted by analyzing RNA-sequencing data using the REPAC pipeline to determine changes in 3’UTR length in treated (HIV-1–infected) versus control (mock-infected) conditions9. 3’ UTRs were designated as shortened if they had compositional fold-change (cFC)<0.25 and −log10 (Adjusted P-Value)<0.05 and as lengthened if they had cFC>0.25 and −log10 (Adjusted P-Value)<0.05.

### PAC-Seq library construction and sequencing

First, 2 µg total RNA was mixed with 1 µl of 100 µM 3’ Illumina_4N_21T_VN primer (5’-GTGACTGGAGTTCAGACGTGTGCTCTTCCGATCTNNNNTTTTTTTTTTTTTTTTTTTTTVN-3’), 2 µl of 5 mM AzNTP (Baseclick, #BCT–25 to −28)/dNTP mixture, at a ratio of 1 to 5, and the total volume was raised to 13 µL by the addition of H_2_O. The mixture was heated at 65°C/5 min and snap-cooled on ice for 3 min. Reverse transcription was performed using SuperScript III (ThermoFisher, #18080093). In brief, 7 µl master mix containing 4 µl 5× Superscript First Strand buffer, 1 µl of 0.1 M DTT, 1 µl RNaseOUT (ThermoFisher, #10777019), and 1 µl SuperScript III were mixed with the RNA sample. Heating was performed in a thermocycler sequentially as follows: 25°C/10 min, 50°C/40 min, and 75°C/15 min, followed by RNase H (ThermoFisher, #AM2293) treatment using 1U per reaction for 37°C/30 min and then 80°C/10 min. The resulting cDNA was purified using Sera-Mag Speedbeads (Cytiva, #65152105050250). Speedbeads working solution was made by washing 1 ml bead slurry twice in 1×TE buffer and then resuspending beads in 50 ml of 1×TE buffer containing 9 g PEG-8000, 1 M NaCl, and 0.05% Tween-20. Speedbeads, 1.8× reaction volume, was mixed with cDNA followed by a 5 min incubation at room temperature. The beads were then pelleted using a magnetic bead collector, and the supernatant was discarded. Two washes with 80% ethanol were performed while the beads were pelleted on the magnet. The beads were then dried, and the cDNA was eluted by resuspending the beads in 22 µl of 50 mM HEPES pH 7.2 for 2 min at room temperature. Then, 20 µl cDNA was mixed with 11 µl Click Mix (2.5 M NaCl, 30% EtOH in H_2_O) and 4 µl of 5 µM UMI-click-adapter (5’-Hexynyl-NNNNNNNNNNNNAGATCGGAAGAGCGTCGTGTAGGGAAAGAGTGT-3’, HPLC purified). To activate the reaction, 4 µl of 50 mM vitamin C and 1 µl of Click Catalyst (4 mM CuSO_4_ in H_2_O and 20 mM THPTA [Baseclick, #BCMI-006] in H_2_O) were pre-mixed before being added into the cDNA solution for 60 min incubation at room temperature in the dark. Clicked cDNA was then purified in the same manner as post-RT purification but eluted in 22 µl of 10 mM Tris pH7.4. Half of the eluted cDNA was subjected to PCR amplification with indexing primers i5 (5’-AATGATACGGCGACCACCGAGATCTACAC[Index]ACACTCTTTCCCTACACGACGC TCTTCCGATC*T-3’) and i7 (5’-CAAGCAGAAGACGGCATACGAGAT[Index]GTGACTGGAGTTCAGACGTGTGCTCTTCCGAT*C-3’). Both primers have a phosphorothioate bond added at the 3’ end (denoted as *) to increase stability. The PCR reaction was conducted in a 50 µl volume, including 25 µl 2× OneTaq Master Mix (NEB, #M0482), 2 µl each of 5 µM i5 and i7, and cDNA in H_2_O. A total of 13 amplification cycles were conducted on 2 µg input RNA. The program was run at 94°C/4 min; 53°C/30 sec; 68°C/10 min; 15× [94°C/30 sec, 53°C/30 sec; 68°C/2 min]; 68°C/5 min; hold at 4°C. The library was then purified and size-selected with Speedbeads. In brief, 45 µl beads (0.9× volume of PCR reaction) were mixed with the reaction to remove larger fragments. Supernatant was collected and mixed with 10 µl beads (0.2× volume of PCR reaction) to remove smaller fragments and primer dimers. Two washes with 100% ethanol were then performed while the beads were on the magnet. Beads were dried before being resuspended in 12 µl of 10 mM Tris pH7.4. The library was then collected and subjected to quality control on Agilent 2200 TapeStation and analysis by Illumina NovaSeq 6000 at the Genomics Research Center at the University of Rochester.

### Pre-processing the raw data derived from PAC-Seq

Samples were sequenced on 10B-300 NovaSeq X-Plus machine with a paired-end 150 bp configuration. For alternative polyadenylation (APA) analysis by PolyA-Miner only forward reads were considered ^44^. Raw fastq reads were subjected to pre-processing steps as described here: Fastp was used to remove the Illumina adapters (using - a option), trim low-quality bases in the start of the read (using -f 4) and to discard reads that are shorter than forty bases (using -l 40) ^53^. The quality trimmed reads were then aligned to the reference genome (GRCh38, version 33 downloaded from GENCODE portal), using bowtie2 version 2.3.5.1 in single end mode. Bowtie2 was run in very sensitive local mode to account for soft clipped regions towards the end of the read. Bowtie2 parameters ^54^: (-p 16 --very-sensitive-local). The alignment files were sorted by genomic coordinates and indexed with samtools version 1.9 ^52^. To address PCR duplicates, Unique Molecular Identifiers (UMI) bar codes were used in the library construction to identify unique transcripts. UMI tools extract version 1.1.5 was used to extract the UMI bar codes from the reads and append to the read names ^55^. Umitools dedup function was used on mapped reads to discard the duplicated reads that share the same UMI ^55^. This will ensure that each unique read is represented once. We then generated the genome wide coverage tracks for these de-duplicated bams in bigWigs format using bamCoverage version 3.5.5 ^56^. Replicates were combined to a single bigWig to be hosted in UCSC Browser to make the gene tracks ^57^. Raw counts were generated from sorted BAMs using featureCounts version v2.0.1 ^58^.

### PolyA-Miner Analysis

PolyA-miner 2022 version was used to quantify alternative polyadenylation ^44^. Parameters used are: -mode bam -s 0 -expNovel 0 -p 20 -ip 50 -a 0.65 -pa_p 0.6 -pa_a 8 -pa_m 4 -gene_min 10 -apa_min 0.05 -t BB. We used bam mode in polyA-miner to process the pre-processed and aligned BAM files (-mode bam). PolyA-miner was run on the following comparisons: HIV-WT infected cells vs Mock cells, HIV-N74D infected cells vs Mock cells, HIV-A77V infected cells vs Mock cells, in A549 cells and T cells and three CPSF6 KO lines vs control cells. Only known polyadenylation sites identified in polyA database were considered for the analysis (-expNovel 0). A polyA site was only considered in the analysis if 60% (-pa_p 0.6) replicates have 8 reads (-pa_a 8) aligned to the site with the third replicate having a minimum of 4 reads (-pa_m 4). A minimum read count of 10 reads per gene is used (-gene_min 10). To avoid sites which were picked due to internal priming, a 20-nucleotide window (-p 20) was set to slide 50 nucleotides (-ip 50) to the downstream of the site. And if in any of the sliding window, there is more than 65% of adenine (-a 0.65), then the site was discarded. All polyA sites retained after mis-priming and other quality filters were included in the downstream APA analysis, regardless of their genomic location (e.g., gene body, UTR exon, or intron). To infer the differential polyA usage variations, beta binomial test was performed using proportions of individual features (polyA signals) to overall gene counts ^58^. For each of the comparisons, polyA-miner generates an output file which includes genes that undergo APA changes. Volcano plots were generated in GraphPad Prism to visualize the changes.

Differential Gene Expression and Gene Ontology Analysis. Gene level counts were calculated as the sum of all the reads mapped to individual polyA signals in each gene. These read counts were normalized and subjected to differential gene expression analysis using DESeq2 version 1.44.0 in R version 4.4.1 (https://www.R-project.org/) Gene ontology analysis was performed using enrichGO function from clusterProfiler 4.12.6 ^59,60^.

## Acknowledgements

We thank the NIH AIDS repository for reagents. C.L., A.Z.D., C.B., B.C., and F.D.-G. are supported by NIH grants AI087390 and AI150455 grants to F.D.-G. We thank Judd Hultquist for helpful discussions.

