## Supplemental data for "HIV-1 Infection Regulates Gene Expression by Altering Alternative Polyadenylation Through CPSF6 and CPSF5 Delocalization"

### Supplementary Tables

**TableS1.** PolyAminer analysis of PAC-Seq 3'UTR APA data from human A549 cells infected with viruses HIV-1-GFP **(A)**, HIV-1-N74D-GFP **(B)** or HIV-1-A77V-GFP **(C)** compared to A549 cells mock-infected, related figure 2.

**TableS2.** PolyAminer analysis of PAC-Seq 3'UTR APA data from Human primary CD4$^+$ T cells infected with viruses HIV-1-GFP **(A)**, HIV-1-N74D-GFP **(B)** or HIV-1-A77V-GFP **(C)** compared to T cells mock-infected, related figure 3.

**TableS3.** PolyAminer analysis of PAC-Seq 3'UTR APA data from human A549- CPSF6-KO cells: CPSF6-KO#B4 **(A)**, CPSF6-KO#B7 **(B)**, and CPSF6-KO#C8 **(C)** compared to A549 parental cells, related figure 5B.

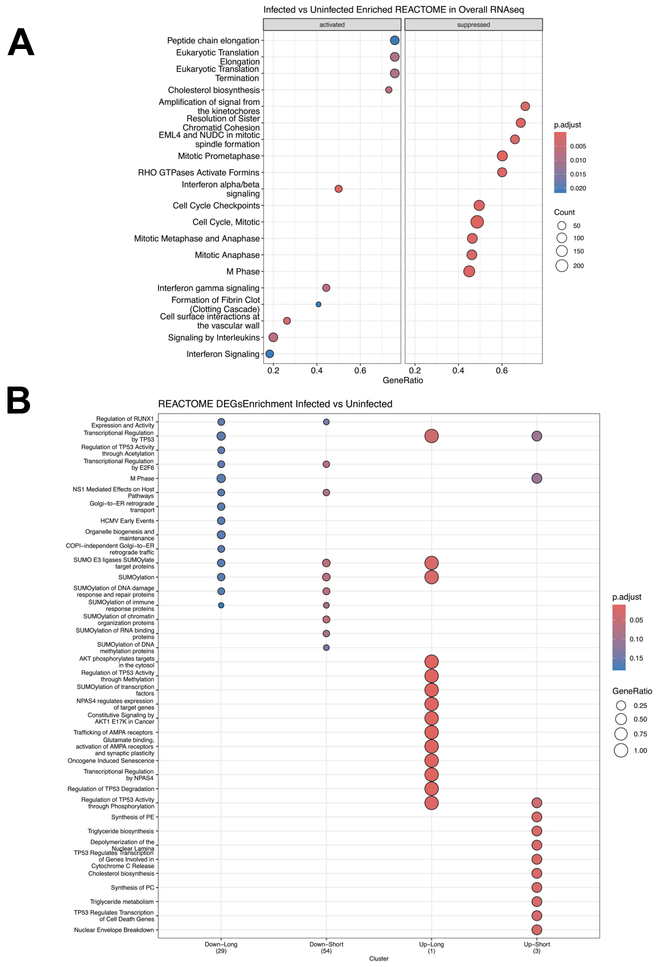


**Figure S1. REACTOME pathway analysis from RNA-seq in A549 cells, related figure 1. (A-B)** Human A549 cells were challenged with HIV-1-GFP viruses at an MOI of ~2 for 24 hours. Total RNA from three infected and three mock-infected samples was sequenced using RNA-seq, as described in the Methods section. **(A)** Pathway enrichment analysis of DEGs. Dot plot shows top 20 enriched REACTOME pathways of the genes that were significantly changed by HIV-1 infection. The size of the dot is based on gene count enriched in the pathway, and the color of the dot shows the pathway enrichment significance (p-value). **(B)** Pathway enrichment analysis of DEGs clustered according to 3'UTR lengths (down-long; down-short; up-long; up-down). The size of the dot is based on GeneRatio enriched in the pathway, and the color of the dot shows the pathway enrichment significance.

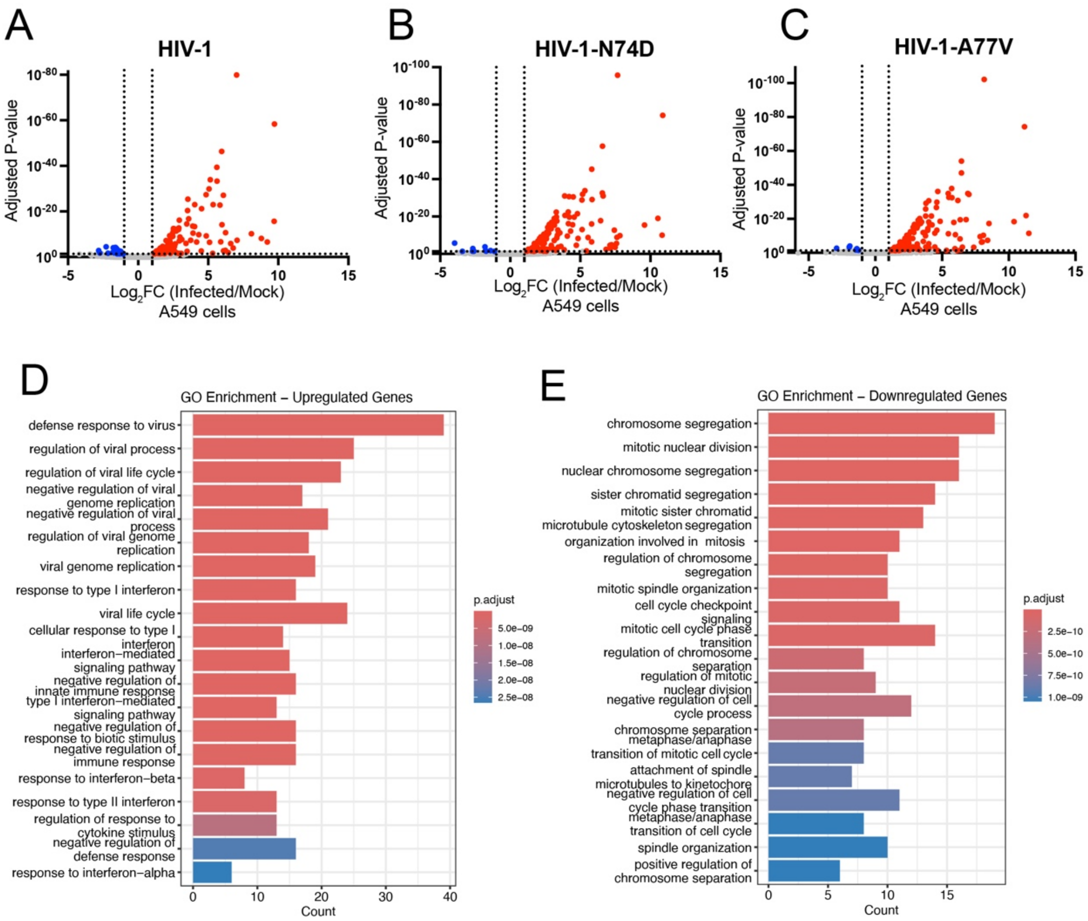


**Figure S2. Gene expression analysis from PAC-seq in A549 cells, related Figure 2.** Human A549 cells were challenged with HIV-1-GFP **(A)**, HIV-1-N74D **(B)** or HIV-1-A77V **(C)** viruses at an MOI of ~2 for 48 hours**.** Total RNA from two infected and two mock-infected samples was sequenced to analyze changes in gene expression, as described in the Methods section. **(A-C)** Volcano plot shows differentially expressed genes in PAC-Seq data. The X-axis represents differences in gene expression as $Log_2$-fold changes. A positive $Log_2$-fold indicates upregulation (red) of the corresponding gene in infected cells, while a negative $Log_2$-fold indicates downregulation (blue). The Y-axis represents the statistical significance of the results, expressed as P-value. **(D-E)** Gene ontology analysis of all genes that were upregulated **(D)** and downregulated **(E)** by HIV-1 infection. The color of the bars represents the P-adjusted value.

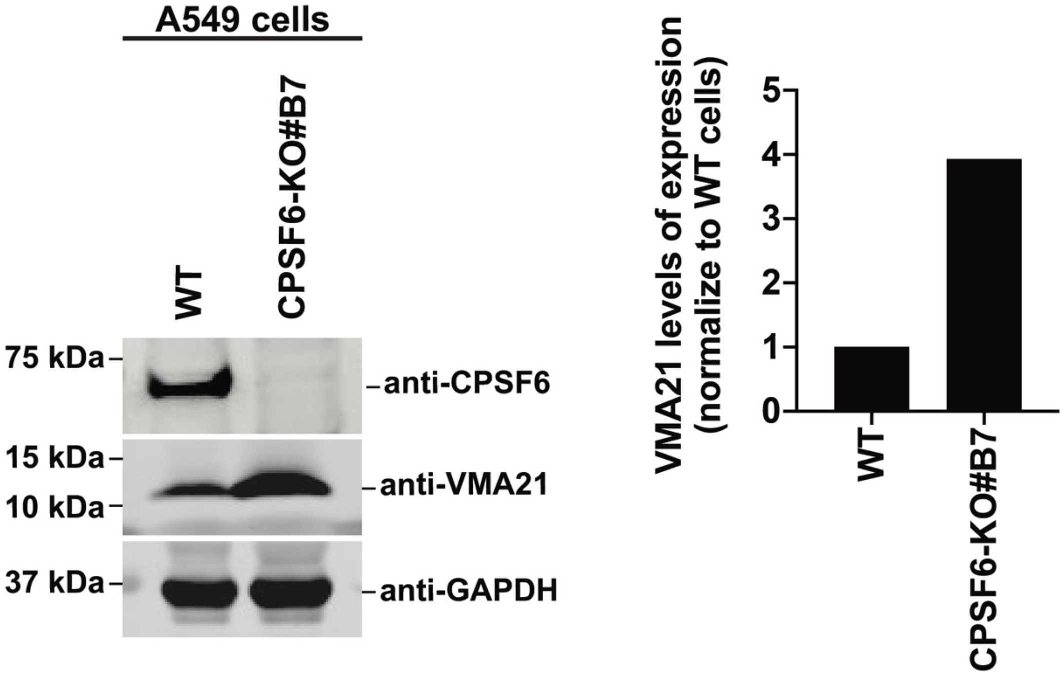


**Figure S3. Loss of CPSF6 expression induces changes in VMA21 protein levels in A549 cells, related Figure 5.** CPSF6-KO#B7 and parental A549 cells were analyzed by Western blotting using anti-VMA21 and anti-CPSF6 antibodies. As loading control, we utilized anti-GAPDH antibodies. Densitometry quantification of the immunoblot signals is shown.
